# Dual orexin/hypocretin receptor antagonism attenuates attentional impairments in an NMDA receptor hypofunction model of schizophrenia

**DOI:** 10.1101/2023.02.05.527043

**Authors:** Eden B. Maness, Sarah A. Blumenthal, Joshua A. Burk

**Author notes:** Please address correspondence to: Eden B. Maness, West Roxbury VA Medical Center, 1400 Veterans of Foreign Wars Parkway, West Roxbury, MA, 02132.

## Abstract

Schizophrenia is a neuropsychiatric condition that is associated with impaired attentional processing and performance. Failure to support increasing attentional load may result, in part, from abnormally overactive basal forebrain projections to the prefrontal cortex, and available antipsychotics often fail to address this issue. Orexin/hypocretin receptors are expressed on corticopetal cholinergic neurons, and their blockade has been shown to decrease the activity of cortical basal forebrain outputs and prefrontal cortical cholinergic neurotransmission. In the present experiment, rats (N = 14) trained in a visual sustained attention task that required discrimination of trials which presented a visual signal from trials during which no signal was presented. Once trained, rats were then co-administered the psychotomimetic N-methyl-D-aspartate (NMDA) receptor antagonist dizocilpine (MK-801: 0 or 0.1 mg/kg, intraperitoneal injections) and the dual orexin receptor antagonist filorexant (MK-6096: 0, 0.1, or 1 mM, intracerebroventricular infusions) prior to task performance across six sessions. Dizocilpine impaired overall accuracy during signal trials, slowed reaction times for correctly-responded trials, and increased the number of omitted trials throughout the task. Dizocilpine-induced increases in signal trial deficits, correct response latencies, and errors of omission were reduced following infusions of the 0.1 mM, but not 1 mM, dose of filorexant. Orexin receptor blockade, perhaps through anticholinergic mechanisms, may improve attentional deficits in a state of NMDA receptor hypofunction.

**Highlights:**

- Schizophrenia is associated with attentional deficits that may stem from abnormally reactive BF projections to the prefrontal cortex
- Orexin receptor antagonists decrease acetylcholine release and reduce prefrontal cortical activity
- The dual orexin receptor antagonist filorexant alleviated impairments of attention following NMDA receptor blockade

## 1. Introduction

Schizophrenia (**SZ**) is a debilitating neurodevelopmental disorder which is characterized by positive symptoms (e.g. hallucinations and delusions), negative symptoms (e.g. anhedonia and avolition), and cognitive symptoms (e.g. attention, learning, and memory impairments). It has been hypothesized that hypofunctional N-methyl-D-aspartate (**NMDA**) receptors expressed on cortical parvalbumin (**PV**)-positive gamma-Aminobutyric acid (**GABA**) interneurons produce chronically disinhibited glutamatergic projections to the midbrain dopamine (**DA**) system and, as a result, evoke sensory processing disturbances associated with greater-than-normal mesolimbic dopaminergic neurotransmission [1, 2, 3]. Extant antipsychotics, all of which antagonize or partially agonize DA-2 receptors, have been widely distributed since the 1950s and provide relief for the hyperdopaminergia-linked positive symptoms [4, 5, 6, 7, 8, 9, 10, 11, 12, 13]. However, in addition to a litany of unpleasant and potentially dangerous side effects associated with a high degree of medicinal non-adherence [14], many patients continue to experience symptoms indicative of a dysfunctional prefrontal cortex (**PFC**), including treatment-resistant deficits of attention [15, 16, 17]. An inability to attend to relevant stimuli and cues is a significant neurocognitive rate-limiting factor and is linked with poor treatment prognosis [18], providing an impetus to investigate novel and alternate pharmacotherapeutic targets.

Controlled attention and the motivated recruitment of attentional effort both rely on the functional integrity of numerous corticopetal, corticofugal, and cortico-cortical projection pathways which together encompass multiple networks devoted to allocating limited cognitive resources towards important stimuli in the external world. One such network is comprised of cortex-innervating basal forebrain (**BF**) cholinergic neurons [19, 20, 21, 22]. Task-relevant release of acetylcholine (**ACh**) in the cortex amplifies the response of pyramidal neurons to incoming sensory information, enhances sensitivity to pertinent cues in the environment, and activates associated response sets [23, 24, 25, 26]. Within the medial PFC, a region that integrates attention and motivation to inform and guide behavior [27, 28, 29, 30], ACh levels rise during behavioral activation and further increase to maintain vigilance in response to attentional challenge [31, 32]. Thus, the stamina and incentive required to sustain attention while filtering out irrelevant stimuli across time largely depend on cholinergic flexibility and adaptability with regards to environmental demands.

Anomalies in the ascending cholinergic system are suggested to be more relevant to the negative and cognitive symptoms than to the positive symptoms in SZ. Corticopetal cholinergic neurons of the BF receive a majority of their inhibitory inputs from GABAergic projections from the nucleus accumbens (**NAcc**) [10, 33, 34, 35]. These projections are suppressed in response to elevated mesolimbic DA neurotransmission in an active disease state, resulting in reduced GABAergic neurotransmission within the BF and ultimately rendering subcortical ACh-generating neurons abnormally and unsustainably reactive in the earliest instances of attentional effort [28, 31, 36, 37]. This BF-PFC dysmorphogenesis is posited to be the primary pathway through which input selection is disrupted and attentional and motivational functioning is impaired as time progresses, especially during periods of environmental distraction [37, 38]. As such, drugs which attenuate BF and PFC overactivity are worthy of exploration as alternative antipsychotic compounds to address ineffective attentional systems associated with psychosis and SZ.

The orexin system, which is also referred to as the hypocretin system, is a widespread neuromodulatory network with cell bodies restricted to the lateral hypothalamus (**LH**) and contiguous perifornical area [39, 40]. The endogenous orexinergic neuropeptides, orexin-A and orexin-B (**OxA** and **OxB**, respectively), are heavily implicated in survival-promoting homeostatic regulatory processes such as appetitive drive and circadian rhythms [39, 40, 41, 42] as well as psychological states including mood, motivation, and cognition [19, 21, 43, 44, 45, 46, 47, 48]. Gq-coupled orexin-1 (**Ox1**) receptors as well as Gq- and Gi/o-linked orexin-2 (**Ox2**) receptors are abundantly co-distributed throughout the BF and PFC, and their activation and enhances interoceptive awareness of internal states and exteroceptive awareness of salient environmental stimuli, potentially through increased prefrontal cortical ACh release [48, 49, 50]. Burgeoning research from Elam and colleagues [51] has shown that in a stress-induced model of psychosis, dual orexin receptor antagonists (**DORAs**) suppress excessive dopaminergic signaling in the ventral tegmental area (**VTA**), dampen the hypersensitivity of DA-producing neurons to stimulant drugs, and normalize behavioral indicators of psychotomimesis (see also [52]). Orexin receptor blockade has also demonstrated anticholinergic tendencies in a disease-free state, as both site-specific and systemic administration of anti-orexinergic ligands dampen cortical ACh release and, at times, impair cortex-dependent cognitive flexibility and attentional performance [49, 53, 54, 55]. DORAs additionally promote GABA release in the BF in addition to other neurotransmitters which decrease cholinergic cell activity through traditionally sleep-promoting pathways, such as adenosine [56]. Therefore, drugs which subdue orexin receptor activity may be effective for improving attentional deficits in conditions associated with BF cholinergic hyperactivity.

To date, there are no studies elucidating the effects of orexin receptor antagonists on attentional performance in SZ, nor has there been research examining DORAs specifically in the context of sustained attention. To address these gaps in the literature, the present experiment measured the effects of intracerebroventricular (**icv**) administration of the DORA filorexant (MK-6096), which has roughly equal affinity for Ox1 and Ox2 receptors [57], on attentional performance of rats following acute intraperitoneal (**ip**) administration of the psychotomimetic NMDA receptor antagonist dizocilpine (MK-801). PV-expressing GABAergic interneurons are particularly sensitive to the effects of subanesthetic concentrations of dizocilpine [58, 59], and similarly to other NMDA receptor antagonists, systemic dizocilpine administration transiently augments cortical ACh levels, induces excitatory-inhibitory imbalance in cognition-associated brain regions, and disrupts attentional performance in a number of paradigms, including a task of sustained attention [3, 60, 61, 62, 63, 64, 65, 66]. Intraventricular infusions of filorexant have translational applicability by assessing the effects of DORAs in widespread brain regions, enabling the opportunity to assess any unexpected deleterious effects from drug actions in brain regions outside of those that are the focus of this experiment. As has been robustly demonstrated, NMDA receptor antagonism through dizocilpine administration was anticipated to negatively impact response accuracy and increase trial omissions, and we hypothesized that the DORA filorexant would alleviate dizocilpine-induced attentional impairments.

## 2. Material and methods

### 2.1. Subjects

A total of 14 adult male Fischer 344/Brown Norway F1 rats (Charles River Laboratories, Wilmington, MA), 12 weeks old upon arrival, were used in the present experiment. Subjects were housed in pairs with a 14-hour light/10-hour dark cycle (lights on 06:00-20:00) in a temperature- and humidity-controlled vivarium, and all behavioral testing occurred 6-7 days per week between 09:00 and 16:00. The rats were allotted *ad libitum* access to rat chow, but water was restricted to 10 minutes a day during testing days and 20 minutes on non-testing days in order to establish water as a salient motivator throughout behavioral training and testing. The protocol for this research was approved by William & Mary’s Institutional Animal Care and Use Committee.

### 2.2. Apparatus

Following the initiation of water restriction, subjects began behavioral testing in one of 14 chambers controlled by MED-PC-V data collection software (Med Associates, Inc., Georgia, VT). Each testing chamber was situated within a sound-attenuating cubicle and consisted of an intelligence panel with one retractable lever on either side of a water access port, a water dipper with a cup which could hold 0.01 ml of water, and a central panel light situated above the water port. A house light was located on the opposite panel of the chamber.

### 2.3. Presurgical behavioral training

Rats were trained in a previously-validated rodent visual sustained attention task (**SAT**) that has previously been shown to be particularly sensitive to BF cholinergic activity and manipulations [67, 68, 69, 70]. The house light remained illuminated throughout the session prior to surgery. The task consisted of three training stages. During the first training stage, both levers were extended throughout testing, and subjects were shaped to press levers using a fixed ratio-1 reinforcement schedule. To prevent the development of a lever bias, if a rat pressed one lever five consecutive times, water access was withheld until a press was made on the other lever. Rats were moved to the next training stage once 120 water rewards were obtained for three successive days. During the second phase of training, which consisted of 100 trials, rats were trained to discriminate between signal trials (1 second illumination of the central panel light) and non-signal trials (no central panel light illumination). After illumination of central panel light (or no illumination), the levers were extended into the chamber. The dipper was raised following a response on one lever on signal trials and the other lever on non-signal trials. Half of the rats received reward access following a response on the left lever on signal trials and following a right lever press on non-signal trials. The correct levers for signal and non-signal trials were reversed for the other half of the animals. Pressing the rewarded lever during a signal trial was considered a hit, and pressing the rewarded lever during a non-signal trial was recorded as a correct rejection. Pressing the incorrect lever in signal trials was considered a miss, and pressing the incorrect lever during a non-signal trial was considered a false alarm. Failure to press either lever after their extension into the testing chamber for 3 seconds was recorded as an omission. Each trial was separated by an inter-trial interval (**ITI**) of 18 seconds. During this training stage, if the rat pressed the incorrect lever or did not press a lever, a correction trial occurred, which was identical to the previous trial. Pressing the incorrect lever for three consecutive correction trials resulted in a forced trial during which only the correct lever was extended into the chamber until a press was made or until 90 seconds elapsed. The central panel light was illuminated if the errors occurred on signal trials. Rats remained in this training stage until accuracy on signal and non-signal trials was at least 70 percent for three consecutive sessions.

The final training stage consisted of 90 trials in each session. There were 45 total signal trials, with an equal number of trials with the 500, 100, and 25 ms signal durations, and 45 non-signal trials (**Figure 1**). Each trial was separated by an ITI of 9 ± 3 seconds. The ITI was shortened and made variable in order to increase the attentional demands of the task. During any given trial, the signal light was either illuminated or not, after which both levers extended for three seconds. Lever pressing was rewarded in the same manner as during the previous training stage. Rats were considered eligible for surgery when they achieved the criteria of 70 percent or higher accuracy in successfully identifying 500 ms signals and at least 70 percent on non-signal trials for three consecutive days.

**Figure 1.**
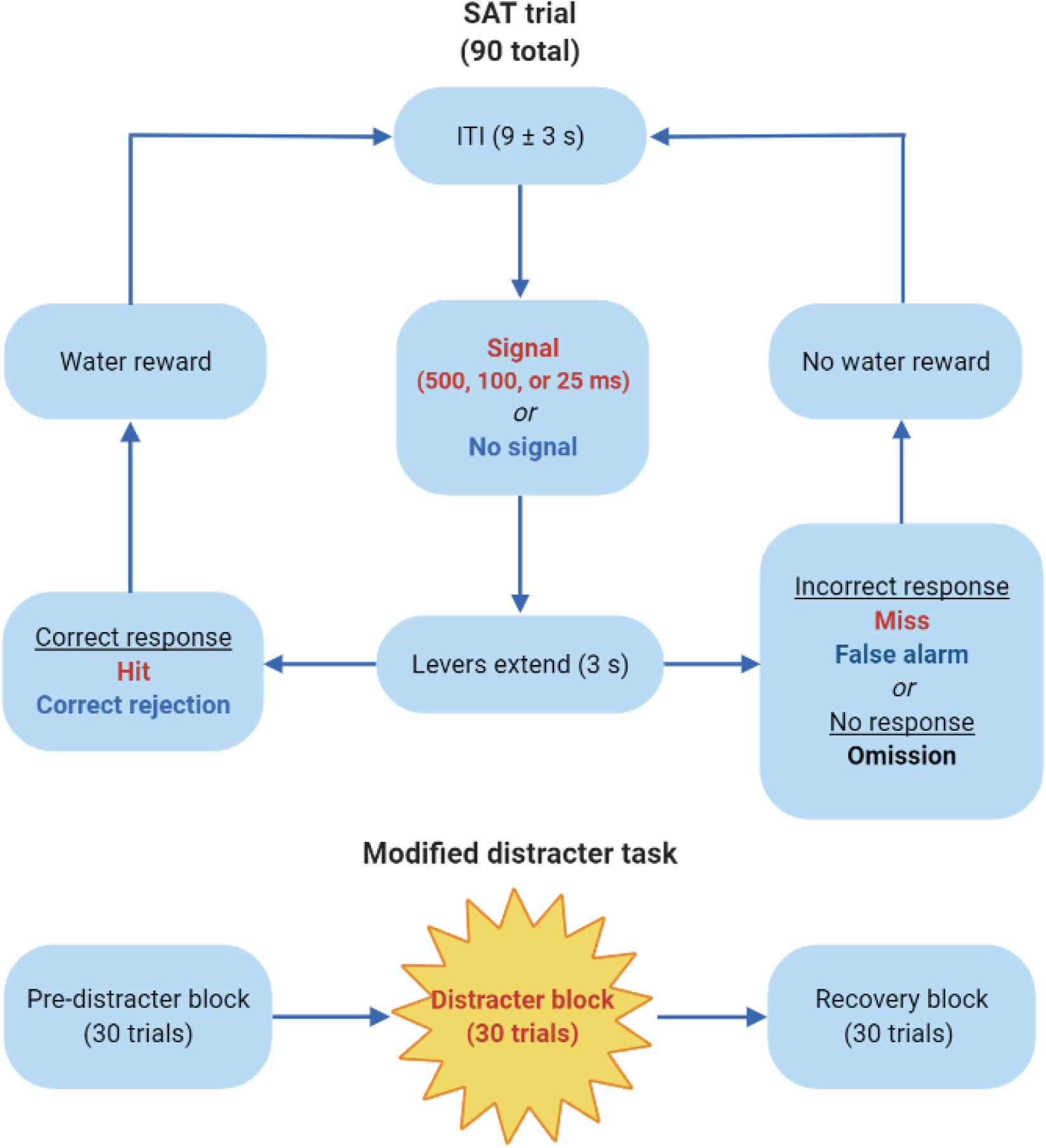
Schematic representation of the sustained attention task (**SAT**) which is comprised of 90 total trials. During a given trial, after an inter-trial interval (**ITI**) between 6 and 12 seconds, a central panel light either flashes (500, 100, or 25 ms) or does not, after which both levers extend into the testing chambers for 3 seconds. If rats press the signal lever at the conclusion of a signal trial, it is recorded as a hit, and they are given a water reward (0.01 ml). Similarly, if the non-signal lever is pressed at the conclusion of a non-signal trial, it is considered a correct rejection, and they are rewarded. However, if they press the non-signal lever following the appearance of the signal (miss) or press the signal lever at the end of a non-signal trial (false alarm), they do not receive a water reward. If they press neither lever within 3 seconds, both levers retract, and it is marked as a trial omission. On drug administration days, the 90 trials are divided into three distinct testing blocks: the pre-distracter block, during which the house light remains illuminated; the distracter block, wherein the house light continuously flashes (0.5 seconds on, 0.5 seconds off); and the recovery block, during which the house light returns to its typical illuminated state.

### 2.4. Surgical procedures

The night prior to surgery, rats were given free access to 2.7 mg/ml of acetaminophen (per os). The following morning, they were anesthetized via an ip injection of 90.0 mg/kg of ketamine and 9.0 mg/kg of xylazine. Upon sedation, rats were shaved around the surgical site, placed in a stereotaxic apparatus (Kopf Instruments, Tujunga, CA), given an incision along the midline, and underwent unilateral icv cannulation surgery wherein one 8.0 mm (22 gauge) guide cannula was implanted 1 mm above either the left or the right lateral ventricle (−0.8 mm anterior-posterior, ±1.6 mm medial-lateral from bregma; -2.5 mm dorsal-ventral from dura; **Figure 2A**). The surgeries were conducted using aseptic conditions, and each cannula was held in place with stainless steel screws and dental cement. After surgery, subjects were given a week-long period of ad libitum water access, with acetaminophen provided for the first three days, after which they resumed water restriction and were reintroduced to the SAT.

**Figure 2.**
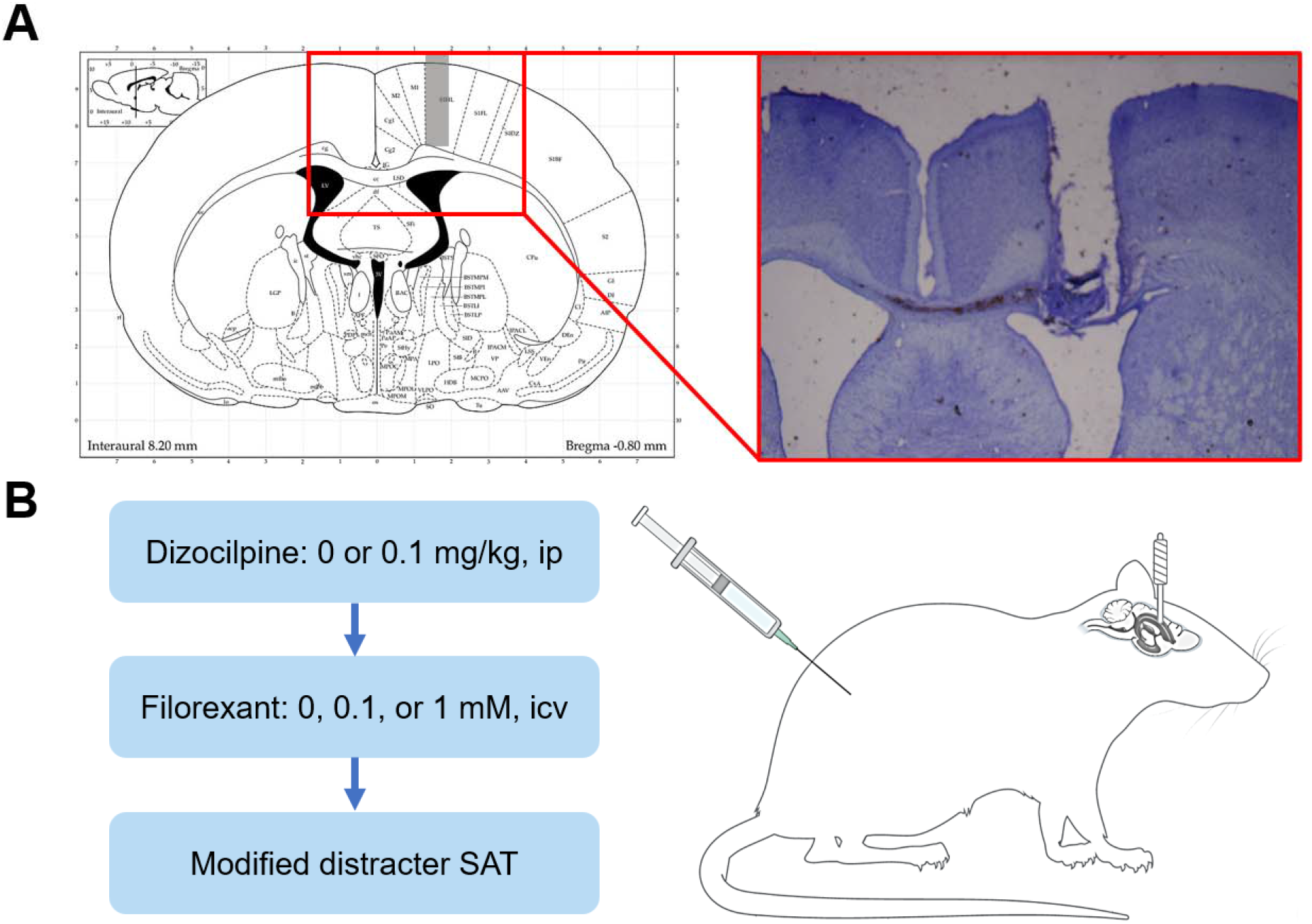
Stereotaxic surgery and drug administration procedures. **2A**. Stereotaxic atlas for intracerebroventricular (**icv**) cannulation and cresyl violet acetate-stained 50 μm section of the lateral ventricles showing cannula tract. Rats underwent unilateral cannulation surgery during which a guide cannula (8 mm, 22 gauge) was placed 1 mm above either the right or left lateral ventricle; internal cannulae extended 1 mm beyond guide cannula into the ventricle. Adjusted from Paxinos & Watson’s *The Rat Brain in Stereotaxic Coordinates*. **2B**. Following surgery, on drug administration days, rats were given intraperitoneal (**ip**) injections of either 0 or 0.1 mg/kg of the N-methyl-D-aspartate receptor antagonist dizocilpine as well as icv infusions of either 0, 0.1, or 1 mM of the dual orexin receptor antagonist filorexant prior to placement in the sustained attention task (**SAT**). Rat image created with BioRender.

### 2.5. Postsurgical behavioral testing

At least one week after baseline performance was re-established, rats were introduced to a modified version of the SAT wherein the house light flashed (0.5 Hz) during the middle set of trials. The 90-trial attention test session was divided into three blocks: the first 30 trials were identical to those of the standard task where with the house light remained illuminated (pre-distracter block), the next 30 trials introduced the flashing house light distracter (distracter block), and the final 30 trials were a return to the standard task with the illuminated house light (recovery block). The rats were exposed to the modified task once before initial drug exposure and on drug administration days; on days when the rats did not receive drugs, they performed in the standard SAT with no distracter.

### 2.6. Drug preparation and administration procedures

Dizocilpine maleate (Tocris Bioscience) was dissolved in sterile saline to create a solution of 0.1 mg/ml that was stored at -40 °C and used within one week of preparation. Stock solutions of 0.1 mM and 1 mM of filorexant (MedChemExpress) in dimethyl sulfoxide (DMSO) were stored at -40 °C and used within one month of preparation.

Rats received a randomized ip injection of either saline or 0.1 mg/kg dizocilpine (at a volume of 1 ml/kg) 30 minutes prior to placement in the chamber. Fifteen minutes before testing, 2.5 μl of either 0 (DMSO vehicle), 0.1, or 1 mM of filorexant was infused at a rate of 1 μl per minute by way of an internal cannula with a 1 mm extension beyond the guide cannula (**Figure 2B**). The internal cannula was connected to a microsyringe on an infusion pump (Harvard Apparatus) via polyethylene tubing. Dizocilpine and filorexant dose combinations were randomized across rats, and a washout period of at least 48 hours separated drug administration sessions.

### 2.7. Histological procedures

Following the final drug administration session, each rat was anesthetized with a ketamine/xylazine cocktail of 100 mg/kg of ketamine and 10 mg/kg of xylazine and transcardially perfused at 300 mmHg with a 10 percent sucrose solution followed by 4 percent paraformaldehyde. Brains were harvested, stored in the 4 percent paraformaldehyde solution, and rinsed with 0.1 M phosphate buffer solution on the day of sectioning. Each brain was sectioned using a vibratome (Thermo Scientific, Microm HM 650V) into 50 μm slices. Sections in the vicinity of the cannulation site were stained with cresyl violet acetate, and guide cannula location was confirmed using an Olympus BX-51 light microscope.

### 2.8. Data analysis

Relative hits were calculated by dividing the number of correct signal identifications by all lever presses at the conclusion of signal trials, and relative correct rejections were determined by dividing the number of correct rejections by the total responses during non-signal trials. These calculations resulted in a number between 0 and 1, with a relative accuracy of 0 indicating 0 percent accuracy and a relative accuracy of 1 indicating 100 percent accuracy. Relative hits, relative correct rejections, and trial omissions were further calculated for pre-distracter, distracter, and recovery blocks. Correct response latency measurements were calculated as the average time (out of 3000 ms) it took rats to press the correct lever following its extension into the testing chamber at the conclusion of a trial, and these were additionally split into blocks for block-specific analyses. Repeated-measures analyses of variance (**ANOVAs**) were used for each measure of accuracy and omissions, all of which were corrected using the Greenhouse-Geisser procedure when required. Interaction effects were further analyzed with one-way repeated-measures ANOVAs, and main effects revealed by these analyses were explored with multiple comparisons of paired-samples *t*-tests and corrected with the Bonferroni pairwise comparison procedure. All statistical analyses were conducted using SPSS Statistics version 24.0. Statistical significance was determined using α = .05. Data are presented as the mean□±□standard error.

## 3. Results

### 3.1. Effects of dizocilpine on SAT performance

Appropriate guide cannula placement was confirmed for all 14 rats included in the behavioral analyses (**Figure 2**). A 2 (dizocilpine: 0 and 0.1 mg/kg) X 3 (block: pre-distracter, distracter, and recovery) X 3 (signal duration: 500, 100, and 25 ms) repeated-measures ANOVA was used to measure the influence of acute NMDA receptor antagonism on signal detection in the SAT. A main effect of dizocilpine (*F*(1,13) = 9.14, *p* = .01, η^2^_*p*_ = .413) revealed that dizocilpine decreased rats’ accuracy in signal trials when compared to saline. There was also a main effect of block (*F*(2,26) = 6.06, *p* = .007, η^2^_*p*_ = .318), with performance being better in the pre-distracter block than the distracter (*t*(13) = 2.27, *p* = .041, *d* = .608) and recovery blocks (*t*(13) = 3.00, *p* = .011, *d* = .796). No interaction between dizocilpine concentration and block suggests that the attentional detriments associated with NMDA receptor blockade are present throughout the testing session rather than during particular periods of testing. Furthermore, a main effect of signal duration (*F*(2,26) = 25.87, *p* < .001, η^2^_*p*_ = .666) revealed an aggregate signal duration-dependent decrease in hit accuracy as signal duration shortened (*p* < .002 between all signal lengths). The signal duration main effect was qualified by a dizocilpine X signal duration interaction (*F*(2,26) = 27.98, *p* < .001, η^2^_*p*_ = .683). Paired-samples *t*-tests at each of the three signal lengths showed that, compared to saline, dizocilpine decreased accuracy following the 500 (*t*(13) = 4.26, *p* = .001, *d* = 1.137) and 100 ms signals (*t*(13) = 2.42, *p* = .031, *d* = .646), but not the 25 ms signal (**Figure 3A**). A dizocilpine X block repeated-measures ANOVA for accuracy in non-signal trials failed to reveal a significant main effect or interaction including dizocilpine as a factor (**Figure 3B**).

**Figure 3.**
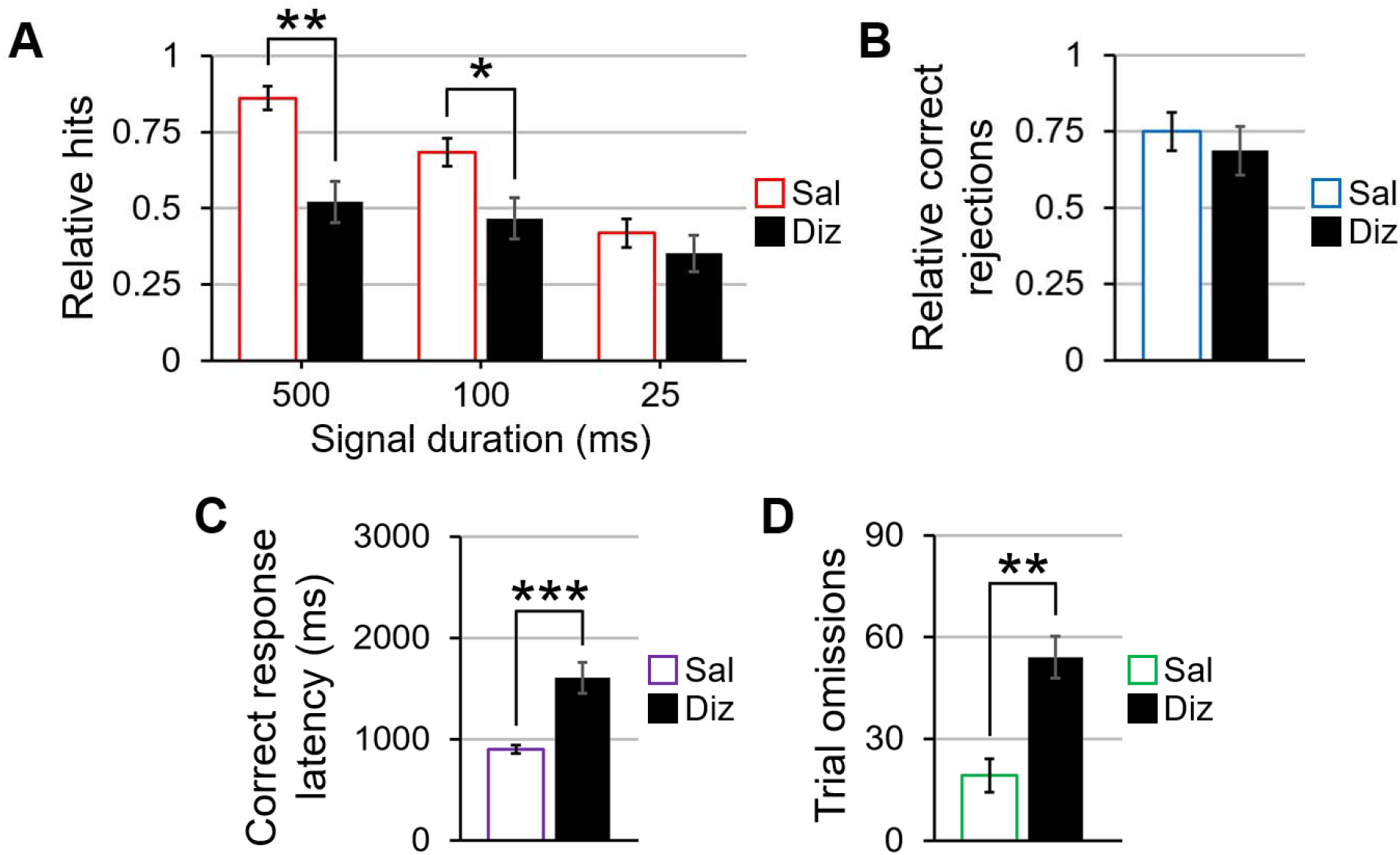
The effects of dizocilpine on signal trials, non-signal trials, reaction times, and omissions in the sustained attention task. **3A**. Compared to when they were given saline, rats given dizocilpine demonstrated a worsened ability to correctly identify the 500 and 100 ms signals, but performance was undisturbed at the 25 ms signal. **3B**. Dizocilpine did not impair performance in non-signal trials. **3C**. Dizocilpine slowed the speed with which rats pressed the correct lever following its extension into the testing chamber at the conclusion of any given trial. **3D**. Dizocilpine significantly increased the number of omitted trials when compared to saline. Error bars are displayed as mean ± SEM. \**p* < .05, \*\**p* < .01, \*\*\**p* < .001

A main effect of dizocilpine in a dizocilpine X block repeated-measures correct response latency ANOVA verified that dizocilpine increased response times when aggregating all trials during which rats responded correctly, *F*(1,13) = 21.93, *p* < .001, η^2^_*p*_ = .628 (**Figure 3C**). Response times also varied by block (*F*(2,26) = 53.14, *p* < .001, η^2^_*p*_ = .803); regardless of whether they were given saline or dizocilpine, rats were slowest to accurately respond in the recovery block when compared to the pre-distracter (*t*(13) = 6.81, *p* < .001, *d* = 1.819) and distracter blocks (*t*(13) = 8.52, *p* < .001, *d* = 2.278), and they were ultimately quickest to respond in the distracter block compared with the pre-distracter block (*t*(13) = 2.82, *p* = .014, *d* = .754). This result suggests that rats responded faster during the flashing house light, but their performance slowed beyond that of pre-distracter speeds once the house light stopped flashing.

For trial omissions, a dizocilpine X block repeated-measures ANOVA revealed a main effect of dizocilpine (*F*(1,13) = 46.72, *p* < .001, η^2^_*p*_ = .782), with NMDA receptor antagonism significantly reducing lever-pressing behavior (**Figure 3D**). The ANOVA also yielded a main effect of block (*F*(2,26) = 20.59, *p* < .001, η^2^_*p*_ = .613), with a block-dependent increase in omissions from pre-distracter to distracter and from distracter to recovery (all *p* < .01). The dizocilpine X block interaction was not statistically significant, indicative of a reduction in on-task behavior regardless of testing period.

### 3.2. Effects of filorexant on SAT performance

To explore the effects of filorexant on attentional performance, a 3 (filorexant: 0, 0.1, and 1 mM) X 3 (block: pre-distracter, distracter, and recovery) X 3 (signal duration: 500, 100, and 25 ms) repeated-measures ANOVA was conducted for signal trial accuracy in the absence of dizocilpine. There was neither a main effect of filorexant nor any interactions between filorexant and other variables of interest on performance in signal and non-signal trials (**Figures 4A** and **4B**, respectively). Similarly, a filorexant X block ANOVA did not reveal any significant effects involving filorexant as a factor. For response latencies in all correctly-responded trials, a filorexant X block repeated-measures ANOVA failed to detect any significant effects of filorexant on reaction times (**Figure 4C**). There was additionally no main or interaction effects of filorexant on omissions following a filorexant X block repeated-measures ANOVA (**Figure 4D**). Therefore, dual orexin receptor antagonism at the doses included in the present experiment does not negatively impact measures of attention in the SAT.

**Figure 4.**
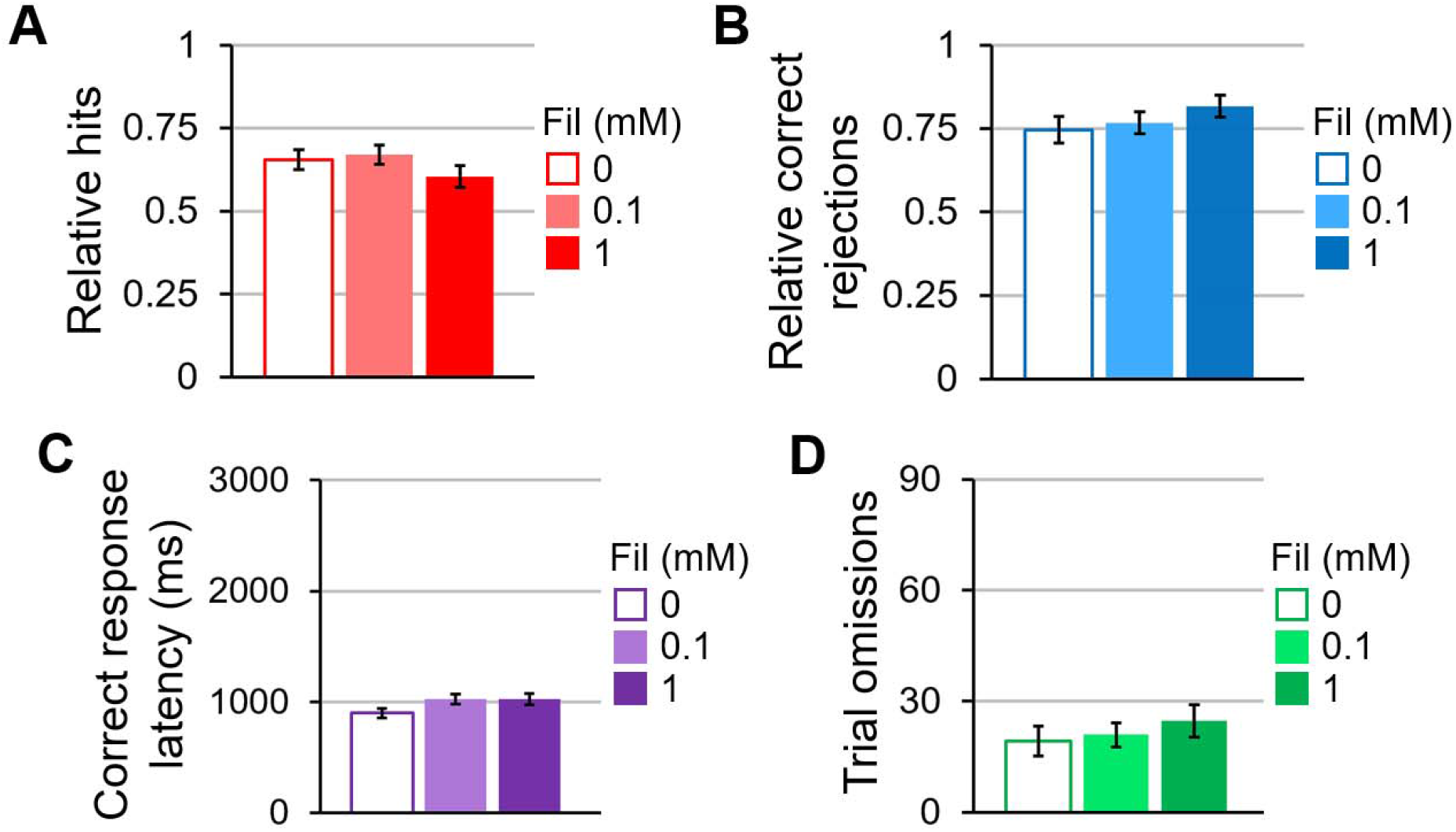
The effects of filorexant on signal trials, non-signal trials, reaction times, and omissions in the sustained attention task. **4A**. Filorexant did not influence accuracy in signal when rats co-administered saline in lieu of dizocilpine. **4B**. Filorexant had no effect on performance in non-signal trials by itself. **4C**. Filorexant did not affect correct response latencies. **4D**. Filorexant did not alter the number of omitted trials across doses on its own. Error bars are displayed as mean ± SEM. \**p* < .05, \*\**p* < .01, \*\*\**p* < .001

### 3.3. Effects of filorexant on dizocilpine-induced SAT impairments

A 2 (dizocilpine: 0 and 0.1 mg/kg) X 3 (filorexant: 0, 0.1, and 1 mM) X 3 (block: pre-distracter, distracter, and recovery) X 3 (signal duration: 500, 100, and 25 ms) repeated-measures ANOVA for signal trial accuracy revealed a main effect of filorexant, *F*(2,26) = 4.42, *p* = .046, η^2^_*p*_ = .254. No significant differences in signal detection were found between the vehicle and the 1 mM filorexant dose, but the 0.1 mM filorexant dose improved signal detection accuracy compared with the vehicle (*t*(13) = 3.13, *p* = .008, d = .836) and 1 mM of filorexant (*t*(13) = 3.54, *p* = .004, *d* = .946; **Figures 5A**). For non-signal trials, a dizocilpine X filorexant X block ANOVA did not yield any significant effects involving dizocilpine or filorexant as factors.

**Figure 5.**
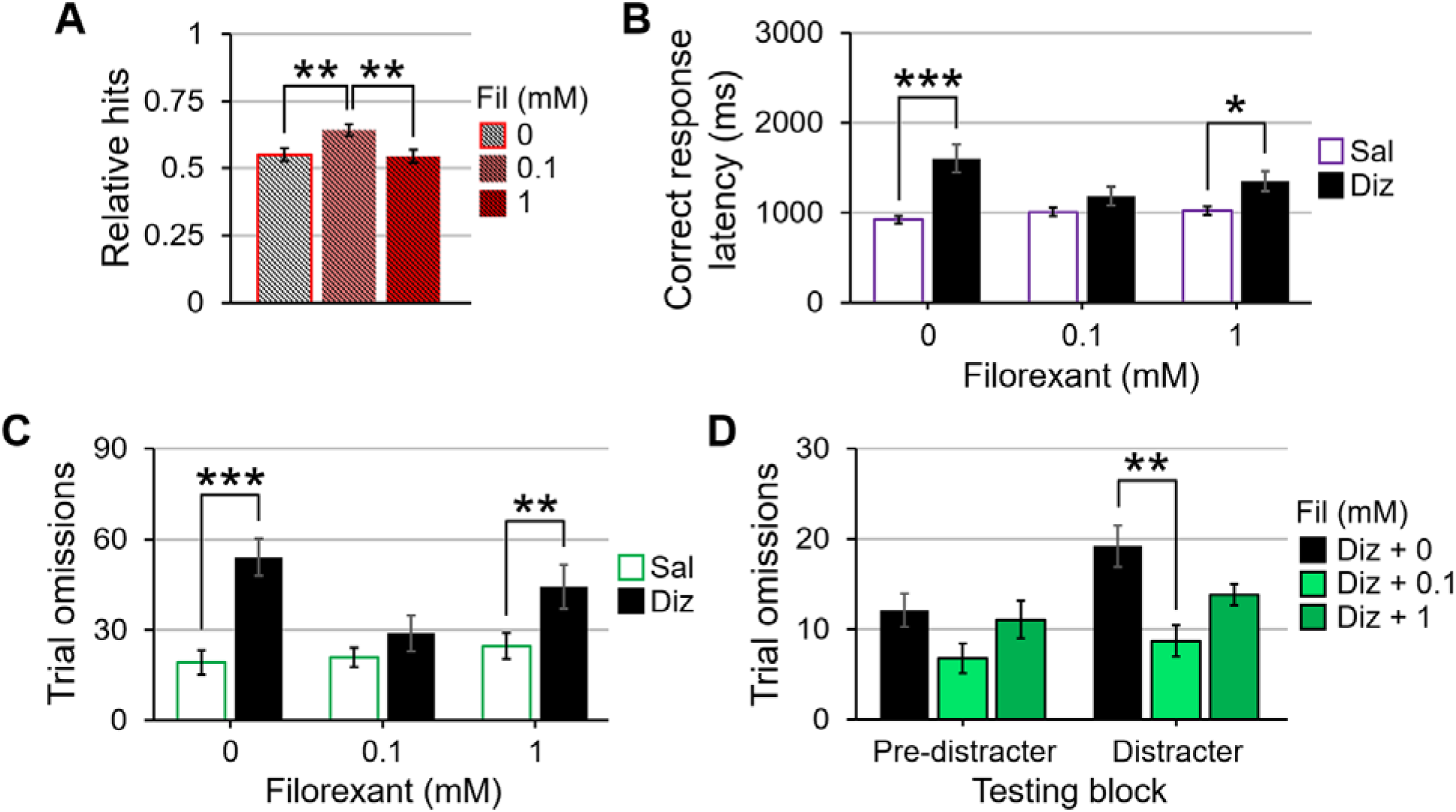
The effects of dizocilpine and filorexant co-administration on signal trial accuracy, reaction times, and omissions in the sustained attention task. **5A**. When averaged across dizocilpine dose, signal trial performance was better when rats were given 0.1 mM of filorexant than when they were given 0 and 1 mM. **5B**. 0.1 mM of filorexant, but not 1 mM, is able to quicken correct response latencies for dizocilpine-administered animals compared to dizocilpine alone. **5C**. When comparing saline- and dizocilpine-injected rats, only the 0.1 mM concentration of filorexant was able to normalize omissions between the two injection conditions. **5D**. When analyses were separated by pre-distracter and distracter blocks, the 0.1 mM dose of filorexant, but not the 1 mM dose, sufficed to significantly improve dizocilpine-induced omissions during the distracter block. Error bars are displayed as mean ± SEM. \**p* < .05, \*\**p* < .01, \*\*\**p* < .001

For reaction times during correctly-responded trials, a dizocilpine X filorexant X block repeated-measures ANOVA revealed a dizocilpine X filorexant interaction (*F*(2,26) = 5.49, *p* = .021, η^2^_*p*_ = .297). Follow-up *t*-tests comparing response speeds when rats were administered saline to when they were administered dizocilpine at each of the three filorexant concentrations showed a slowing of reaction times when rats were co-administered dizocilpine and 0 mM (*t*(13) = 4.68, *p* < .001, *d* = 1.251) as well as with 1 mM doses of filorexant (*t*(13) = 2.44, *p* = .03, *d* = .652); however, no such difference was detected between saline- and dizocilpine-injected rats when they were given infusions of 0.1 mM of filorexant (**Figure 5B**). This dizocilpine X filorexant two-way interaction was further qualified by a dizocilpine X filorexant X block three-way interaction (*F*(4,52) = 5.22, *p* = .007, η^2^_*p*_ = .287), indicating that the influence of filorexant on dizocilpine-associated slowing of reaction times partly relied on testing period. Dizocilpine X filorexant repeated-measures ANOVAs for each block revealed a main effect of dizocilpine in each of the three testing periods (all *p* < .005) where rats took longer to respond following NMDA receptor blockade regardless of the introduction of filorexant to the analyses. While no block had a main effect of filorexant, there were dizocilpine X filorexant interactions in the pre-distracter (*F*(2,26) = 4.43, *p* = .039, η^2^_*p*_ = .254) and recovery periods (*F*(2,26) = 7.56, *p* = .002, η^2^_*p*_ = .654). Dizocilpine concentration *t*-tests for each filorexant dose in these two blocks showed that rats were slower to press the correct lever when they were given dizocilpine than when they were given saline in the absence of filorexant (*p* < .003 for both blocks). However, both 0.1 and 1 mM concentrations of filorexant improved reaction times of dizocilpine-administered rats to that of baseline (both *p* > .05 when compared to infusions of vehicle in both blocks), highlighting a beneficial influence of orexin receptor antagonism in quickening reaction times following NMDA receptor blockade before and after the flashing house light.

A dizocilpine X filorexant X block repeated-measures omissions ANOVA revealed a main effect of filorexant (*F*(2,26) = 5.17, *p* = .025, η^2^_*p*_ = .293), a filorexant X dizocilpine interaction (*F*(2,26) = 12.14, *p* < .001, η^2^_*p*_ = .432), and a dizocilpine X filorexant X block interaction (*F*(4,52) = 3.90, *p* = .008, η^2^_*p*_ = .231) on the number of omitted trials. Together, these findings suggest that the effects of filorexant on dizocilpine-induced omissions vary depending on testing period. When averaged across block, dizocilpine dose paired-samples *t*-tests at each of the three filorexant doses reveal that, while dizocilpine-administered rats omitted more when co-administered infusions of vehicle (*t*(13) = 6.84, *p* < .001, *d* = 1.827) and 1 mM of filorexant (*t*(13) = 3.36, *p* = .005, *d* = .898), the 0.1 mM concentration equalized trial omissions between the two injection conditions (**Figure 5C**). When divided into pre-distracter, distracter, and recovery blocks, dizocilpine X filorexant ANOVAs uncovered dizocilpine-filorexant interactions prior to (*F*(2,26) = 4.04, *p* = .03, η^2^_*p*_ = .237) and during the flashing distracter (*F*(2,26) = 7.29, *p* = .003, η^2^_*p*_ = .359). For rats given 0.1 mg/kg of dizocilpine, a main effect of filorexant was present only during the flashing distracter (*F*(2,26) = 6.76, *p* = .004, η^2^_*p*_ = .359). Multiple comparisons *t*-tests juxtaposing the three filorexant doses revealed that omissions were statistically similar between 0 and 1 mM doses of filorexant for dizocilpine-administered rats, but for those co-administered dizocilpine and 0.1 mM of filorexant, the number of omissions was significantly reduced from when they were given dizocilpine alone (**Figure 5D**; *t*(13) = 3.34, *p* = .005, *d* = .893). Altogether, these omission analyses highlight a robust ameliorative effect of the lower filorexant concentration in the SAT.

## 4. Discussion

The findings from the present experiment demonstrate that acute systemic administration of dizocilpine, a psychotomimetic often employed to emulate the NMDA receptor hypofunction observed in SZ, decreased accuracy in signal trials, increased response latency during correct trials, and increased trial omissions in the SAT. Because performance in non-signal trials remained intact, it suggests that rats were still able to respond based upon the task rules when dizocilpine was administered, and decreased signal detection accuracy is not due to a side or lever bias. Regardless of trial type, dizocilpine augmented response latency for trials during which rats responded at the correct lever, a well-established finding that has been replicated in other animal studies using single-dose administration of NMDA receptor antagonists in similar paradigms of signal detection, such as the five-choice serial reaction time task (**5CSRTT**) [61, 71, 72]. Lastly, dizocilpine increased trial omissions in all task blocks compared to saline, suggesting that motivation to respond in the SAT may also be negatively impacted in this model of SZ. Motoric impairments cannot be fully discounted, as locomotion was not directly quantified in this study, though no serious aberrations of movement were observed prior to placement in and following removal from the task chamber.

To date, this experiment is the first to explore the influence of dual orexin receptor antagonism on sustained attentional performance concurrently with a pharmacological model of SZ. Filorexant was hypothesized to ameliorate decreases in accuracy as well as increases in correct response latency and omissions when co-administered with an NMDA receptor antagonist. Consistent with this hypothesis, filorexant attenuated deficits in hit accuracy for rats administered dizocilpine, though the effects are modest here. Widespread blockade of orexin receptors - especially those located on cells which are known to proliferate frontocortical ACh efflux - may have slightly improved signal detection though the reduction of dizocilpine-produced excitatory-inhibitory disturbances in sustained attention-relevant cortical areas. Additionally, increases in correct response latencies induced by dizocilpine were abated following co-administration of the low dose of filorexant in both signal and non-signal trials. This finding is clinically applicable, as some previous research using dizocilpine and other NMDA receptor antagonist models of SZ have shown that antipsychotic pretreatment either does not affect or can exacerbate increases in correct response and reward retrieval latencies [66, 73].

In addition to improving attentional accuracy, intracranial infusions of filorexant reduced the number of omitted trials resulting from dizocilpine exposure. Based on our behavioral findings, it can be surmised that dual orexin receptor antagonism may have attenuated abnormally elevated prefrontal cortical stimulatory neurotransmission that precluded appropriate adjustment to increasing attentional load. In doing so, the incentive to maintain performance in this pharmacological model of SZ was partly restored. Interestingly, the 0.1 mM concentration of filorexant, but not the 1 mM concentration, was sufficient to restore in-task responding to levels observed with saline injections, with the high dose producing a similarly elevated number of omissions as dizocilpine alone. The neurobiological mechanisms underlying the adverse outcomes regarding in-task responsiveness following the combination of dizocilpine and 1 mM of filorexant observed in this experiment remain unclear. However, the effects of orexin receptor blockade on the pursuit of reinforcement are heavily influenced by the effort-to-reward ratio [74] - that is, high-effort, but not low-effort, response capacity is differentially impacted by orexinergic antagonists. Because dizocilpine made consistent performance in the SAT more effortful, the exertion required to maintain performance across trials may have exceeded the drive to attain the water reward. While this motivational deactivation was reversed by a lower degree of orexin receptor antagonism, findings from the present experiment suggest that higher DORA concentrations may be insufficient in improving task engagement in this model of acute psychosis. It can therefore be speculated that lower concentrations of anti-orexinergic drugs do not necessarily impact incentivized performance or motoric functioning at a concentration that benefitted rats in the SAT in the context of NMDA receptor hypofunction.

In addition to putatively anticholinergic effects in the BF, dual orexin receptor antagonism may have indirectly addressed dizocilpine-induced dysfunctions of signal-driven input selection and cortical vigilance networks stemming via various non-cholinergic loci in the mesocorticolimbic system, including PV+ GABAergic neurons of the BF. These fast-spiking neurons - which have recently been shown to quicken reaction times in a rodent psychomotor vigilance task when optogenetically stimulated [75, 76] - are depolarized by orexins and are themselves sufficient to stimulate cortical arousal when exposed to OxA in animals with lesions to corticopetal cholinergic projections [43, 77]. Thus, frontocortical activation may be suppressed when blocking orexin receptors expressed on these neurons as well as orexin receptors located on cortical glutamatergic outputs to the BF, which synapse exclusively with these PV+ GABAergic neurons [78, 79].

Moreover, because muted GABAergic efferents from the NAcc to BF cholinergic neurons are hypothesized to contribute to the attentional deficits in SZ [80], orexin receptor antagonism may mitigate the prevalence of SZ-linked behavioral correlates by modulating mesolimbic DA synthesis. This idea is supported by findings from Elam et al. [51] showing that the DORA suvorexant, the Ox1 receptor-specific selective orexin receptor antagonist (**SORA**) SB-334867, and the Ox2 receptor-targeting SORA EMPA reduced VTA DA neuron population activity in a rodent model of stress-induced psychosis, and the former two reversed dizocilpine-induced hyperlocomotion through presumed antidopaminergic mechanisms of action. Neurons in the cortex containing both Ox1 and Ox2 receptor mRNA also project to and synapse with A10 DA cells of the nucleus paranigralis subdivision of the VTA and the shell of the NAcc [81, 82, 83, 84], so it is possible that DORAs may lessen BF hyper-reactivity through the inhibition of these orexin-sensitive corticofugal innervations.

It is also feasible that filorexant has attentionally-beneficial effects outside of the BF and dopaminergic midbrain, including through reciprocal connections between the LH and the PFC [85, 86], midline-intralaminar thalamic relay neurons [87, 88], and norepinephrinergic neurons of the locus coeruleus [45, 53, 89, 90, 91]. Site-specific administration can provide better insight as to which vigilance-relevant brain circuits respond to and benefit the most from orexin receptor-targeting ligands in models of NMDA receptor hypofunction.

In the absence of dizocilpine, neither dose of filorexant impacted accuracy or correct reaction times in the SAT, nor did they diminish lever-pressing behavior. This is a clinically-relevant finding, as ideal drug candidates to treat attentional impairment in SZ would not worsen cognition on their own. As has been discussed, functionality of Ox1 receptors is required for homeostasis-driven pursuit of food [84, 92, 93, 94], water [95, 96], and drug reinforcers [97, 98, 99], and Ox2 receptor activity plays a critical role in the instigation and maintenance of consciousness, behavioral arousal, and general awareness [100, 101]. The composite findings of this experiment suggest that, in the absence of other psychoactive pharmacological compounds, non-specific blockade of both orexin receptor subtypes neither interfered with motivation to perform the SAT nor induced lethargy that precluded adequate responding. This is corroborated by research from Gentile and colleagues [102] showing that orexin receptor blockade is able to lessen cocaine-induced premature responses in the 5CSRTT without de-incentivizing reward-contingent attentional performance. It also offers further support for the aforementioned idea that orexin receptor antagonists may primarily demotivate the pursuit of reinforcement during periods of augmented effort, such as when they are administered in tandem with other compounds known to disrupt focus-based performance [74, 103, 104].

Though the presented findings highlight a putatively therapeutic benefit of DORAs for the treatment of attention-relevant deficits in the context of NMDA receptor hypofunction, the gap in the literature can be further addressed outside of the scope of the present experiment. For example, the inclusion of female rats is important to parse any sex-specific effects of the drugs used in the study; though less is known about sex differences in response to orexin receptor antagonists, it has been shown that female rats may be more sensitive to the influence of dizocilpine and other NMDA receptor-blocking compounds [105, 106]. Site-specific, rather than intraventricular, administration of orexin receptor-targeting agents can offer insight regarding which specific brain regions and their associated behaviors are most responsive to anti-orexinergic compounds. The employment of SORAs can additionally elucidate the unique roles of the Ox1 and Ox2 receptor subtypes in influencing attentional outcomes in this model of acute psychosis. Histochemical techniques, such as c-Fos staining and immunofluorescence, can reveal neurobiological effects of DORA exposure that, together with behavioral interpretations, offer a more complete understanding of cellular and behavioral outcomes of orexinergic manipulations.

## 5. Conclusions

In an acute NMDA receptor antagonism model of SZ, dual orexin receptor blockade was able to alleviate dizocilpine-induced alterations of signal trial performance, reaction time, and trial omissions in a test of sustained attention. In particular, the 0.1 mM concentration of filorexant improved response accuracy, restored correct response latency, and lessened the number of omitted trials. Besides improving reaction times in the pre- and post-distracter periods, the 1 mM dose failed to demonstrate many of the same ameliorative qualities, suggesting that a higher degree of orexin receptor inhibition fails to improve upon or incentivize task performance for rats when they are co-administered dizocilpine. This experiment is the first to introduce orexin receptors as a potential novel pharmacotherapeutic target to treat the sustained attentional and perhaps motivational deficits associated with psychosis. The employment of SORAs to research the role of individual orexin receptor subtypes on brain physiology and behavior in this model is a logical next step towards a more complete picture of the therapeutic potential of orexin receptors for the treatment of attentional impairments in SZ.

## Acknowledgments

This work was supported by NIH R01 AG050518 (JAB). The authors would like to thank Angela Goyal, Bella Salas, Derek Griffin, Faith Reid, Grace Smith, Gwyneth Pudner, Lauren Burnette, Nickash Sivakumar, Paige Little, Rahul Patel, Reema Patel, Saurav Pattanayak, and Stacy Pitcairn for their technical assistance.

## References

[1] Homayoun, H. & Moghaddam, B. (2007). NMDA Receptor Hypofunction Produces Opposite Effects on Prefrontal Cortex Interneurons and Pyramidal Neurons. The Journal of Neuroscience, 27(43), 11496–11500. https://doi.org/10.1523/JNEUROSCI.2213-07.2007

[2] Hyde, T. M. & Crook, J. M. (2001). Cholinergic systems and schizophrenia: primary pathology or epiphenomena? Journal of Chemical Neuroanatomy, 22(1-2), 53–63. https://doi.org/10.1016/S0891-0618(01)00101-6

[3] Jentsch, J. D. & Roth, R. H. (1999). The Neuropsychopharmacology of Phencyclidine: From NMDA Receptor Hypofunction to the Dopamine Hypothesis of Schizophrenia. Neuropsychopharmacology, 20(3), 201–225. https://doi.org/10.1016/S0893-133X(98)00060-8

[4] Davis, K. L., Kahn, R. S., Ko, G., & Davidson, M. (1991). Dopamine in schizophrenia: a review and reconceptualization. American Journal of Psychiatry, 148(11), 1474–1486. https://doi.org/10.1176/ajp.148.11.1474

[5] Jentsch, J. D., Redmond, D. E., Elsworth, J. D., Taylor, J. R., Youngren, K. D., & Roth, R. H. (1997a). Enduring cognitive deficits and cortical dopamine dysfunction in monkeys after long-term administration of phencyclidine. Science, 277(5328), 953–955. https://doi.org/10.1126/science.277.5328.953

[6] Johnstone, E. C., Crow, T. J., Frith, C. D., Carney, M. W., & Price, J. S. (1978). Mechanism of the antipsychotic effect in the treatment of acute schizophrenia. Lancet, 1(8069), 845–851. https://doi.org/10.1016/s0140-6736(78)90193-9

[7] Kay, S. R., Fiszbein, A., & Opler, L. A. (1987). The positive and negative syndrome scale (PANSS) for schizophrenia. Schizophrenia Bulletin, 13(2), 261–276. https://doi.org/10.1093/schbul/13.2.261

[8] Lawler, C. P., Prioleau, C., Lewis, M. M., Mak, C., Jiang, D., Schetz, J. A., Gonzales, A. M., Sibley, D. R., & Mailman, R. B. (1999). Interactions of the novel antipsychotic aripiprazole (OPC-14597) with dopamine and serotonin receptor subtypes. Neuropsychopharmacology, 20(6), 612–627. https://doi.org/10.1016/S0893-133X(98)00099-2

[9] Lieberman, J. A. (2004). Dopamine partial agonists: a new class of antipsychotic. CNS Drugs, 18(4), 251–267. https://doi.org/10.2165/00023210-200418040-00005

[10] Sarter, M. & Bruno, J. P. (1998). Cortical acetylcholine, reality distortion, schizophrenia, and lewy body dementia: too much or too little cortical acetylcholine? Brain and Cognition, 38(3), 297–316. https://doi.org/10.1006/brcg.1998.1035

[11] Stahl, S. M. (2008). Stahl’s Essential Psychopharmacology, fourth ed. Cambridge, UK; New York: Cambridge University Press.

[12] Tamminga, C. A., Buchanan, R. W., & Gold, J. M. (1998). The role of negative symptoms and cognitive dysfunction in schizophrenia outcome. International Clinical Psychopharmacology, 13(Suppl 3), S21–S26. https://doi.org/10.1097/00004850-199803003-00004

[13] Tandon, R. (2011). Antipsychotics in the treatment of schizophrenia: an overview. Journal of Clinical Psychiatry, 72, 4–8. https://doi.org/10.4088/JCP.10075su1.01

[14] Zhang, J. & Malhotra, A. K. (2011). Pharmacogenetics and antipsychotics: therapeutic efficacy and side effects prediction. Expert Opinion on Drug Metabolism & Toxicology, 7(1), 9–37. https://doi.org/10.1517/17425255.2011.532787

[15] Lowe, P., Krivoy, A., Porffy, L., Henriksdottir, E., Eromona, W., & Shergill, S. S. (2018). When the drugs don’t work: treatment-resistant schizophre nia, serotonin and serendipity. Therapeutic Advances in Psychopharmacology, 8(1), 63–70. https://doi.org/10.1177/2045125317737003

[16] Reilly, J. L., Harris, M. S. H., Khine, T. T., Keshavan, M. S., & Sweeney, J. A. (2007). Antipsychotic Drugs Exacerbate Impairment on a Working Memory Task in First-Episode Schizophrenia. Biological Psychiatry, 62(7), 818–821. https://doi.org/10.1016/j.biopsych.2006.10.031

[17] van Os, J. & Kapur, S. (2009). Schizophrenia. The Lancet, 374(9690), 635–645. https://doi.org/10.1016/S0140-6736(09)60995-8

[18] Green, M. F., Kern, R. S., & Heaton, R. K. (2004). Longitudinal studies of cognition and functional outcome in schizophrenia: implications for MATRICS. Schizophrenia Research, 72(1), 41–51. https://doi.org/10.1016/j.schres.2004.09.009

[19] Burk, J. A., Blumenthal, S. A., & Maness, E. B. (2018). Neuropharmacology of attention. European Journal of Pharmacology, 835, 162–168. https://doi.org/10.1016/j.ejphar.2018.08.008

[20] Eggermann, E., Serafin, M., Bayer, L., Machard, D., Saint-Mleux, B., Jones, B. E., & Mühlethaler, M. (2001). Orexins/hypocretins excite basal forebrain cholinergic neurones. Neuroscience, 108(2), 177–181. https://doi.org/10.1016/s0306-4522(01)00512-7

[21] Fadel, J. & Burk, J. A. (2010). Orexin/hypocretin modulation of the basal forebrain cholinergic system: role in attention. Brain Research, 1314, 112–123. https://doi.org/10.1016/j.brainres.2009.08.046

[22] Sarter, M. & Bruno, J. P. (1999). Cortical cholinergic inputs mediating arousal, attentional processing and dreaming: differential afferent regulation of the basal forebrain by telencephalic and brainstem afferents. Neuroscience, 95(4), 933–952. https://doi.org/10.1016/S0306-4522(99)00487-X

[23] Dickinson, J. A., Kew, J. N. C., & Wonnacott, S. (2008). Presynaptic α7- and β2-Containing Nicotinic Acetylcholine Receptors Modulate Excitatory Amino Acid Release from Rat Prefrontal Cortex Nerve Terminals via Distinct Cellular Mechanisms. Molecular Pharmacology, 74(2), 348–359. https://doi.org/10.1124/mol.108.046623

[24] Gill, T. M., Sarter, M., & Givens, B. (2000). Sustained Visual Attention Performance-Associated Prefrontal Neuronal Activity: Evidence for Cholinergic Modulation. The Journal of Neuroscience, 20(12), 4745–4757. https://doi.org/10.1523/JNEUROSCI.20-12-04745.2000

[25] Higley, M. J. & Picciotto, M. R. (2014). Neuromodulation by acetylcholine: examples from schizophrenia and depression. Current Opinion in Neurobiology, 29, 88–95. https://doi.org/10.1016/j.conb.2014.06.004

[26] Perry, E., Walker, M., Grace, J., & Perry, R. (1999). Acetylcholine in mind: a neurotransmitter correlate of consciousness? Trends in Neurosciences, 22(6), 273–280. https://doi.org/10.1016/S0166-2236(98)01361-7

[27] Dalley, J. W., Cardinal, R. N., & Robbins, T. W. (2004). Prefrontal executive and cognitive functions in rodents: neural and neurochemical substrates. Neuroscience & Biobehavioral Reviews, 28(7), 771–784. https://doi.org/10.1016/j.neubiorev.2004.09.006

[28] Lustig, C. & Sarter, M. (2015). Attention and the Cholinergic System: Relevance to Schizophrenia. In T. W. Robbins & B. J. Sahakian (Eds.), Translational Neuropsychopharmacology (Vol. 28, pp. 327–362). Springer International Publishing. https://doi.org/10.1007/7854_2015_5009

[29] Small, D. M., Gitelman, D. R., Gregory, M. D., Nobre, A. C., Parrish, T. B., & Mesulam, M.-M. (2003). The posterior cingulate and medial prefrontal cortex mediate the anticipatory allocation of spatial attention. NeuroImage, 18(3), 633–641. https://doi.org/10.1016/S1053-8119(02)00012-5

[30] Watanabe, M. & Sakagami, M. (2007). Integration of Cognitive and Motivational Context Information in the Primate Prefrontal Cortex. Cerebral Cortex, 17(suppl 1), i101–i109. https://doi.org/10.1093/cercor/bhm067

[31] Kozak, R., Bruno, J. P., & Sarter, M. (2006). Augmented Prefrontal Acetylcholine Release during Challenged Attentional Performance. Cerebral Cortex, 16(1), 9–17. https://doi.org/10.1093/cercor/bhi079

[32] St. Peters, M., Demeter, E., Lustig, C., Bruno, J. P., & Sarter, M. (2011). Enhanced Control of Attention by Stimulating Mesolimbic-Corticopetal Cholinergic Circuitry. The Journal of Neuroscience, 31(26), 9760–9771. https://doi.org/10.1523/JNEUROSCI.1902-11.2011

[33] Salgado, S. & Kaplitt, M.G. (2015). The nucleus accumbens: a comprehensive review. Stereotactic and Functional Neurosurgery, 93(2), 75–93. https://doi.org/10.1159/000368279

[34] Záborszky, L., Cullinan, W. E., & Braun, A. (1991). Afferents to Basal Forebrain Cholinergic Projection Neurons: An Update. In T. C. Napier, P. W. Kalivas, & I. Hanin (Eds.), The Basal Forebrain (Vol. 295, pp. 43–100). Springer US. https://doi.org/10.1007/978-1-4757-0145-6_2

[35] Záborszky, L. & Cullinan, W. E. (1992). Projections from the nucleus accumbens to cholinergic neurons of the ventral pallidum: a correlated light and electron microscopic double-immunolabeling study in rat. Brain Research, 570(1-2), 92–101. https://doi.org/10.1016/0006-8993(92)90568-T

[36] Manoach, D. S. (2003). Prefrontal cortex dysfunction during working memory performance in schizophrenia: reconciling discrepant findings. Schizophrenia Research, 60(2-3), 285–298. https://doi.org/10.1016/S0920-9964(02)00294-3

[37] Sarter, M., Hasselmo, M. E., Bruno, J. P., & Givens, B. (2005). Unraveling the attentional functions of cortical cholinergic inputs: interactions between signal-driven and cognitive modulation of signal detection. Brain Research Reviews, 48(1), 98–111. https://doi.org/10.1016/j.brainresrev.2004.08.006

[38] Demeter, E., Guthrie, S. K., Taylor, S. F., Sarter, M., & Lustig, C. (2013). Increased distractor vulnerability but preserved vigilance in patients with schizophrenia: Evidence from a translational Sustained Attention Task. Schizophrenia Research, 144(1-3), 136–141. https://doi.org/10.1016/j.schres.2013.01.003

[39] Peyron, C., Tighe, D. K., van den Pol, A. N., de Lecea, L., Heller, H. C., Sutcliffe, J. G., & Kilduff, T. S. (1998). Neurons containing hypocretin (orexin) project to multiple neuronal systems. The Journal of Neuroscience, 18(23), 9996–10015. https://doi.org/10.1523/JNEUROSCI.18-23-09996.1998

[40] Sakurai, T., Amemiya, A., Ishii, M., Chemelli, R. M., Tanaka, H., Williams, S. C., Richardson, J. A., Kozlowski, G. P., Wilson, S., Arch, J. R., Buckingham, R. E., Haynes, A. C., Carr, S. A., Annan, R. S., McNulty, D. E., Liu, W. S., Terrett, J. A., Elshourbagy, N. A., Bergsma, D. J., & Yanagisawa, M. (1998). Orexins and orexin receptors: a family of hypothalamic neuropeptides and G protein-coupled receptors that regulate feeding behavior. Cell, 92(4), 573–585. https://doi.org/10.1016/s0092-8674(00)80949-6

[41] Dube, M. G., Kalra, S. P., & Kalra, P. S. (1999). Food intake elicited by central administration of orexins/hypocretins: identification of hypothalamic sites of action. Brain Research, 842(2), 473–477. https://doi.org/10.1016/s0006-8993(99)01824-7

[42] Tsujino, N. & Sakurai, T. (2009). Orexin/hypocretin: a neuropeptide at the interface of sleep, energy homeostasis, and reward system. Pharmacological Reviews, 61(2), 162–176. https://doi.org/10.1124/pr.109.001321

[43] Arrigoni, E., Mochizuki, T., & Scammell, T. E. (2010). Activation of the basal forebrain by the orexin/hypocretin neurones. Acta Physiologica (Oxf), 198(3), 223–235. https://doi.org/10.1111/j.1748-1716.2009.02036.x

[44] Eriksson, K. S., Sergeeva, O. A., Haas, H. L., & Selbach, O. (2010). Orexins/hypocretins and aminergic systems. Acta Physioliga (Oxf), 198(3), 263–275. https://doi.org/10.1111/j.1748-1716.2009.02015.x

[45] Mahler, S. V., Moorman, D. E., Smith, R. J., James, M. H., & Aston-Jones, G. (2014). Motivational activation: a unifying hypothesis of orexin/hypocretin function. Nature Neuroscience, 17(10), 1298–1303. https://doi.org/10.1038/nn.3810

[46] Selbach, O., Doreulee, N., Bohla, C., Eriksson, K. S., Sergeeva, O. A., Poelchen, W., Brown, R. E., & Haas, H. L. (2004). Orexins/hypocretins cause sharp wave- and theta-related synaptic plasticity in the hippocampus via glutamatergic, gabaergic, noradrenergic, and cholinergic signaling. Neuroscience, 127(2), 519–528. https://doi.org/10.1016/j.neuroscience.2004.05.012

[47] Turner, D. C., Clark, L., Dowson, J., Robbins, T. W., & Sahakian, B. J. (2004). Modafinil improves cognition and response inhibition in adult attention-deficit/hyperactivity disorder. Biological Psychiatry, 55(10), 1031–1040. https://doi.org/10.1016/j.biopsych.2004.02.008

[48] Zajo, K. N., Fadel, J. R., & Burk, J. A. (2015). Orexin A-induced enhancement of attentional processing in rats: Role of basal forebrain neurons. Psychopharmacology (Berl), 233(4), 639–647. https://doi.org/10.1007/s00213-015-4139-z

[49] Dong, H., Fukuda, S., Murata, E., Zhu, Z., & Higuchi, T. (2006). Orexins Increase Cortical Acetylcholine Release and Electroencephalographic Activation through Orexin-1 Receptor in the Rat Basal Forebrain during Isoflurane Anesthesia. Anesthesiology, 104(5), 1023–1032. https://doi.org/10.1097/00000542-200605000-00019

[50] Fadel, J. & Deutch, A. Y. (2002). Anatomical substrates of orexin-dopamine interactions: lateral hypothalamic projections to the ventral tegmental area. Neuroscience, 111(2), 379–387. https://doi.org/10.1016/s0306-4522(02)00017-9

[51] Elam, H. B., Perez, S. M., Donegan, J. J., & Lodge, D. J. (2021). Orexin receptor antagonists reverse aberrant dopamine neuron activity and related behaviors in a rodent model of stress-induced psychosis. Translational Psychiatry, 11(1), 114. https://doi.org/10.1038/s41398-021-01235-8

[52] Perez, S. M. & Lodge, D. J. (2021). Orexin modulation of VTA dopamine neuron activity: relevance to schizophrenia. International Journal of Neuropsychopharmacology, 24(4), 344–353. https://doi.org/10.1093/ijnp/pyaa080

[53] Boschen, K. E., Fadel, J. R., & Burk, J. A. (2009). Systemic and intrabasalis administration of the orexin-1 receptor antagonist, SB-334867, disrupts attentional performance in rats. Psychopharmacology, 206(2), 205–213. https://doi.org/10.1007/s00213-009-1596-2

[54] Frederick-Duus, D., Guyton, M. F., & Fadel, J. (2007). Food-elicited increases in cortical acetylcholine release require orexin transmission. Neuroscience, 149(3), 499–507. https://doi.org/10.1016/j.neuroscience.2007.07.061

[55] Piantadosi, P. T., Holmes, A., Roberts, B. M., & Bailey, A. M. (2015). Orexin receptor activity in the basal forebrain alters performance on an olfactory discrimination task. Brain Research, 1594, 215–222. https://doi.org/10.1016/j.brainres.2014.10.041

[56] Vazquez-DeRose, J., Schwartz, M. D., Nguyen, A. T., Warrier, D. R., Gulati, S., Mathew, T. K., Neylan, T. C., & Kilduff, T. S. (2016). Hypocretin/orexin antagonism enhances sleep-related adenosine and GABA neurotransmission in rat basal forebrain. Brain Structure and Function, 221(2), 923–940. https://doi.org/10.1007/s00429-014-0946-y

[57] Winrow, C. J., Gotter, A. L., Cox, C. D., Tannenbaum, P. L., Garson, S. L., Doran, S. M., Breslin, M. J., Schreier, J. D., Fox, S. V., Harrell, C. M., Stevens, J., Reiss D. R., Cui, D., Coleman, P. J., & Renger, J. J. (2012). Pharmacological characterization of MK-6096 - a dual orexin receptor antagonist for insomnia. Neuropharmacology, 62(2), 978–987. https://doi.org/10.1016/j.neuropharm.2011.10.003

[58] Bygrave, A. M., Masiulis, S., Nicholson, E., Berkemann, M., Barkus, C., Sprengel, R., Harrison, P. J., Kullmann, D. M., Bannerman, D. M., & Kätzel, D. (2016). Knockout of NMDA-receptors from parvalbumin interneurons sensitizes to schizophrenia-related deficits induced by MK-801. Translational Psychiatry, 6(4), e778–e778. https://doi.org/10.1038/tp.2016.44

[59] Xi, D., Zhang, W., Wang, H.-X., Stradtman, G. G., & Gao, W.-J. (2009). Dizocilpine (MK-801) induces distinct changes of N-methyl-d-aspartic acid receptor subunits in parvalbumin-containing interneurons in young adult rat prefrontal cortex. The International Journal of Neuropsychopharmacology, 12(10), 1395. https://doi.org/10.1017/S146114570900042X

[60] Acquas, E., Wilson, C., & Fibiger, H. C. (1998). Pharmacology of sensory stimulation-evoked increases in frontal cortical acetylcholine release. Neuroscience, 85(1), 73–83. https://doi.org/10.1016/s0306-4522(97)00546-0

[61] Amitai, N. & Markou, A. (2010). Disruption of performance in the five-choice serial reaction time task induced by administration of N-methyl-D-aspartate receptor antagonists: relevance to cognitive dysfunction in schizophrenia. Biological Psychiatry, 68(1), 5–16. https://doi.org/10.1016/j.biopsych.2010.03.004

[62] Coyle, J. T. (2006). Glutamate and schizophrenia: beyond the dopamine hypothesis. Cell Molecular Neurobiology, 26(4-6), 365–384. https://doi.org/10.1007/s10571-006-9062-8

[63] Giovannini, M., Mutolo, D., Bianchi, L., Michelassi, A., & Pepeu, G. (1994). NMDA receptor antagonists decrease GABA outflow from the septum and increase acetylcholine outflow from the hippocampus: a microdialysis study. The Journal of Neuroscience, 14(3), 1358–1365. https://doi.org/10.1523/JNEUROSCI.14-03-01358.1994

[64] Howe, W. M. & Burk, J. A. (2007). Dizocilpine-induced accuracy deficits in a visual signal detection task are not present following D-cycloserine administration in rats. European Journal of Pharmacology, 577(1-3), 87–90. https://doi.org/10.1016/j.ejphar.2007.08.037

[65] Manahan-Vaughan, D., von Haebler, D., Winter, C., Juckel, G., & Heinemann, U. (2007). A single application of MK801 causes symptoms of acute psychosis, deficits in spatial memory, and impairments of synaptic plasticity in rats. Hippocampus, 18(2), 125–134. https://doi.org/10.1002/hipo.20367

[66] Paine, T. A. & Carlezon, W. A., Jr. (2009). Effects of antipsychotic drugs on MK-801-induced attention and motivational deficits in rats. Neuropharmacology, 56(4), 788–797. https://doi.org/10.1016/j.neuropharm.2009.01.004

[67] Arnold, H. M., Burk, J. A., Hodgson, E. M., Sarter, M., & Bruno, J. P. (2002). Differential cortical acetylcholine release in rats performing a sustained attention task versus behavioral control tasks that do not explicitly tax attention. Neuroscience, 114(2), 451–460. https://doi.org/10.1016/s0306-4522(02)00292-0

[68] Bushnell, P. J. & Strupp, B. J. (2009). Assessing Attention in Rodents. In J. J. Buccafusco (Ed.), Methods of Behavior Analysis in Neuroscience (2nd ed.). CRC Press/Taylor & Francis.

[69] McGaughy, J. & Sarter, M. (1995). Behavioral vigilance in rats: task validation and effects of age, amphetamine, and benzodiazepine receptor ligands. Psychopharmacology, 117(3), 340–357. https://doi.org/10.1007/BF02246109

[70] Sarter, M., Givens, B., & Bruno, J. P. (2001). The cognitive neuroscience of sustained attention: where top-down meets bottom-up. Brain Research. Brain Research Reviews, 35(2), 146–160. https://doi.org/10.1016/s0165-0173(01)00044-3

[71] Nikiforuk, A. & Popik, P. (2014). The effects of acute and repeated administration of ketamine on attentional performance in the five-choice serial reaction time task in rats. European Neuropsychopharmacology, 24(8), 1381–1393. https://doi.org/10.1016/j.euroneuro.2014.04.007

[72] Smith, J. W., Gastambide, F., Gilmour, G., Dix, S., Foss, J., Lloyd, K., Malik, N., & Tricklebank, M. (2011). A comparison of the effects of ketamine and phencyclidine with other antagonists of the NMDA receptor in rodent assays of attention and working memory. Psychopharmacology, 217(2), 255–269. https://doi.org/10.1007/s00213-011-2277-5

[73] Amitai, N., Semenova, S., & Markou, A. (2007). Cognitive-disruptive effects of the psychotomimetic phencyclidine and attenuation by atypical antipsychotic medications in rats. Psychopharmacology, 193(4), 521–537. https://doi.org/10.1007/s00213-007-0808-x

[74] Shaw, J. K., Black, E. M., Zhang, Y., & España, R. A. (2019). Hypocretin Receptor 1 Regulation of Dopamine Neurotransmission and Motivated Behavior. In The Orexin/Hypocretin System (pp. 99–120). Elsevier. https://doi.org/10.1016/B978-0-12-813751-2.00005-X

[75] Maness, E.B., Burk, J.A., Schiffino, F.L., McKenna, J.T., Strecker, R.E., & McCoy, J.G. (2022). Role of the locus coeruleus and basal forebrain in arousal and attention. Brain Research Bulletin, 188, 47–58. https://doi.org/10.1016/j.brainresbull.2022.07.014

[76] Schiffino, F. L., McNally, J. M., Brown, R. E., & Strecker, R. E. (2021). Basal forebrain parvalbumin neurons modulate vigilant attention. bioRxiv. https://doi.org/10.1101/2021.04.19.440515

[77] Blanco-Centurion, C., Gerashchenko, D., & Shiromani, P. J. (2007). Effects of Saporin-Induced Lesions of Three Arousal Populations on Daily Levels of Sleep and Wake. The Journal of Neuroscience, 27(51), 14041–14048. https://doi.org/10.1523/JNEUROSCI.3217-07.2007

[78] Sarter, M. & Bruno, J. P. (2002). The neglected constituent of the basal forebrain corticopetal projection system: GABAergic projections: Cortical GABA afferents. European Journal of Neuroscience, 15(12), 1867–1873. https://doi.org/10.1046/j.1460-9568.2002.02004.x

[79] Záborszky, L., Gaykema, R. P., Swanson, D. J., & Cullinan, W. E. (1997). Cortical input to the basal forebrain. Neuroscience, 79(4), 1051–1078. https://doi.org/10.1016/S0306-4522(97)00049-3

[80] Sarter, M., Bruno, J. P., & Turchi, J. (1999). Basal Forebrain Afferent Projections Modulating Cortical Acetylcholine, Attention, and Implications for Neuropsychiatric Disorders. Annals of the New York Academy of Sciences, 877(1), 368–382. https://doi.org/10.1111/j.1749-6632.1999.tb09277.x

[81] Carr, D. B. & Sesack, S. R. (2000). Projections from the Rat Prefrontal Cortex to the Ventral Tegmental Area: Target Specificity in the Synaptic Associations with Mesoaccumbens and Mesocortical Neurons. The Journal of Neuroscience, 20(10), 3864–3873. https://doi.org/10.1523/JNEUROSCI.20-10-03864.2000

[82] Murase, S., Grenhoff, J., Chouvet, G., Gonon, F. G., & Svensson, T. H. (1993). Prefrontal cortex regulates burst firing and transmitter release in rat mesolimbic dopamine neurons studied in vivo. Neuroscience Letters, 157(1), 53–56. https://doi.org/10.1016/0304-3940(93)90641-w

[83] Sesack, S. R. & Pickel, V. M. (1992). Prefrontal cortical efferents in the rat synapse on unlabeled neuronal targets of catecholamine terminals in the nucleus accumbens septi and on dopamine neurons in the ventral tegmental area. The Journal of Comparative Neurology, 320(2), 145–160. https://doi.org/10.1002/cne.903200202

[84] Thorpe, A. J. & Kotz, C. M. (2005). Orexin A in the nucleus accumbens stimulates feeding and locomotor activity. Brain Research, 1050(1-2), 156–162. https://doi.org/10.1016/j.brainres.2005.05.045

[85] Gabbott, P. L. A., Warner, T. A., Jays, P. R. L., Salway, P., & Busby, S. J. (2005). Prefrontal cortex in the rat: Projections to subcortical autonomic, motor, and limbic centers. The Journal of Comparative Neurology, 492(2), 145–177. https://doi.org/10.1002/cne.20738

[86] Kita, H. & Oomura, Y. (1981). Reciprocal connections between the lateral hypothalamus and the frontal cortex in the rat: Electrophysiological and anatomical observations. Brain Research, 213(1), 1–16. https://doi.org/10.1016/0006-8993(81)91244-0

[87] Hay, Y. A., Andjelic, S., Badr, S., & Lambolez, B. (2015). Orexin-dependent activation of layer VIb enhances cortical network activity and integration of non-specific thalamocortical inputs. Brain Structure and Function, 220(6), 3497–3512. https://doi.org/10.1007/s00429-014-0869-7

[88] Lambe, E. K., Liu, R.-J., & Aghajanian, G. K. (2006). Schizophrenia, Hypocretin (Orexin), and the Thalamocortical Activating System. Schizophrenia Bulletin, 33(6), 1284–1290. https://doi.org/10.1093/schbul/sbm088

[89] Aston-Jones, G., Rajkowski, J., & Cohen, J. (1999). Role of locus coeruleus in attention and behavioral flexibility. Biological Psychiatry, 46(9), 1309–1320. https://doi.org/10.1016/S0006-3223(99)00140-7

[90] Foote, S. L., Berridge, C. W., Adams, L. M., & Pineda, J. A. (1991). Electrophysiological evidence for the involvement of the locus coeruleus in alerting, orienting, and attending. In Progress in Brain Research (Vol. 88, pp. 521–532). Elsevier. https://doi.org/10.1016/S0079-6123(08)63831-5

[91] Trivedi, P., Yu, H., MacNeil, D. J., Van der Ploeg, L. H. T., & Guan, X.-M. (1998). Distribution of orexin receptor mRNA in the rat brain. FEBS Letters, 438(1-2), 71–75. https://doi.org/10.1016/S0014-5793(98)01266-6

[92] Bingham, M. J., Cai, J., & Deehan, M. R. (2006). Eating, sleeping and rewarding: orexin receptors and their antagonists. Current Opinion in Drug Discovery & Development, 9(5), 551–559.

[93] Choi, D. L., Davis, J. F., Fitzgerald, M. E., & Benoit, S. C. (2010). The role of orexin-A in food motivation, reward-based feeding behavior and food-induced neuronal activation in rats. Neuroscience, 167(1), 11–20. https://doi.org/10.1016/j.neuroscience.2010.02.002

[94] Sharf, R., Sarhan, M., Brayton, C. E., Guarnieri, D. J., Taylor, J. R., & DiLeone, R. J. (2010). Orexin Signaling Via the Orexin 1 Receptor Mediates Operant Responding for Food Reinforcement. Biological Psychiatry, 67(8), 753–760. https://doi.org/10.1016/j.biopsych.2009.12.035

[95] Hurley, S. W. & Johnson, A. K. (2014). The role of the lateral hypothalamus and orexin in ingestive behavior: a model for the translation of past experience and sensed deficits into motivated behaviors. Frontiers in Systems Neuroscience, 8. https://doi.org/10.3389/fnsys.2014.00216

[96] Kunii, K., Yamanaka, A., Nambu, T., Matsuzaki, I., Goto, K., & Sakurai, T. (1999). Orexins/hypocretins regulate drinking behaviour. Brain Research, 842(1), 256–261. https://doi.org/10.1016/S0006-8993(99)01884-3

[97] Hutcheson, D. M., Quarta, D., Halbout, B., Rigal, A., Valerio, E., & Heidbreder, C. (2011). Orexin-1 receptor antagonist SB-334867 reduces the acquisition and expression of cocaine-conditioned reinforcement and the expression of amphetamine-conditioned reward. Behavioural Pharmacology, 22(2), 173–181. https://doi.org/10.1097/FBP.0b013e328343d761

[98] James, M. H., Mahler, S. V., Moorman, D. E., & Aston-Jones, G. (2016). A Decade of Orexin/Hypocretin and Addiction: Where Are We Now? In A. J. Lawrence & L. de Lecea (Eds.), Behavioral Neuroscience of Orexin/Hypocretin (Vol. 33, pp. 247–281). Springer International Publishing. https://doi.org/10.1007/7854_2016_57

[99] Moorman, D. E., James, M. H., Kilroy, E. A., & Aston-Jones, G. (2017). Orexin/hypocretin-1 receptor antagonism reduces ethanol self-administration and reinstatement selectively in highly-motivated rats. Brain Research, 1654, 34–42. https://doi.org/10.1016/j.brainres.2016.10.018

[100] Mieda, M., Tsujino, N., & Sakurai, T. (2013). Differential Roles of Orexin Receptors in the Regulation of Sleep/Wakefulness. Frontiers in Endocrinology, 4. https://doi.org/10.3389/fendo.2013.00057

[101] Sasaki, K., Suzuki, M., Mieda, M., Tsujino, N., Roth, B., & Sakurai, T. (2011). Pharmacogenetic Modulation of Orexin Neurons Alters Sleep/Wakefulness States in Mice. PLoS ONE, 6(5), e20360. https://doi.org/10.1371/journal.pone.0020360

[102] Gentile, T. A., Simmons, S. J., Watson, M. N., Connelly, K. L., Brailoiu, E., Zhang, Y., & Muschamp, J. W. (2018). Effects of Suvorexant, a Dual Orexin/Hypocretin Receptor Antagonist, on Impulsive Behavior Associated with Cocaine. Neuropsychopharmacology, 43(5), 1001–1009. https://doi.org/10.1038/npp.2017.158

[103] Brodnik, Z. D., Bernstein, D. L., Prince, C. D., & España, R. A. (2015). Hypocretin receptor 1 blockade preferentially reduces high effort responding for cocaine without promoting sleep. Behavioural Brain Research, 291, 377–384. https://doi.org/10.1016/j.bbr.2015.05.051

[104] España, R. A., Oleson, E. B., Locke, J. L., Brookshire, B. R., Roberts, D. C. S., & Jones, S. R. (2010). The hypocretin-orexin system regulates cocaine self-administration via actions on the mesolimbic dopamine system. European Journal of Neuroscience, 31(2), 336–348. https://doi.org/10.1111/j.1460-9568.2009.07065.x

[105] Andiné, P., Widermark, N., Axelsson, R., Nyberg, G., Olofsson, U., Mõrtensson, E., & Sandberg, M. (1999). Characterization of MK-801-induced behavior as a putative rat model of psychosis. The Journal of Pharmacology and Experimental Therapeutics, 290(3), 1393–1408.

[106] Hur, G.-H., Son, W.-C., Shin, S., Kang, J.-K., & Kim, Y.-B. (1999). Sex differences in dizocilpine (MK-801) neurotoxicity in rats. Environmental Toxicology and Pharmacology, 7(2), 143–146. https://doi.org/10.1016/S1382-6689(99)00003-4

